# Sparks fade with distance: The effect of electric field distribution on global motion perception using different tES techniques

**DOI:** 10.1101/2025.01.20.633831

**Authors:** Andrea Pavan, Filippo Ghin, Adriano Contillo, Sibel Akyuz, Gianluca Campana

**Affiliations:** University of Bologna, Department of Psychology, Viale Berti Pichat, 5, 40127 Bologna, Italy; University of Lincoln, School of Psychology, Brayford Wharf East, Lincoln, LN5 7AY, United Kingdom; Cognitive Neurophysiology, Department of Child and Adolescent Psychiatry, Faculty of Medicine of the TU Dresden, Fetscherstraße 74, Schubertstraße 42, 01309, Dresden, Germany; Elettra-Sincrotrone Trieste S.C.p.A, 34149, Trieste, Italy; University of Bergamo, Department of Human and Social Sciences, Bergamo, Italy; University of Padova, Department of General Psychology, Via Venezia 8, 35131 Padova, Italy; Human Inspired Technology Research Centre, University of Padova, Via Luzzati 4, 35121 Padova, Italy

**Author notes:** **Corresponding Author** Andrea Pavan University of Bologna Department of Psychology Viale Berti Pichat, 5, 40127, Bologna, Italy.

**Keywords:** global motion, transcranial electrical stimulation, electric field, extracephalic montage

## Abstract

Previous evidence has shown that high frequency transcranial random noise stimulation (hf-tRNS) decreases motion coherence thresholds when a cephalic montage (i.e., return over Cz) is used. Extracephalic montages have also been employed to modulate behavioral performance, eliminating stimulation of regions under the return electrode. In this study, we examined the effects of different transcranial electrical stimulation (tES) protocols on visual motion discrimination, placing the return electrode on the ipsilateral arm. We assessed the impact of electrode localization using hf-tRNS, anodal, cathodal transcranial direct current stimulation (tDCS), and Sham stimulation over hMT^+^, a brain region involved in global motion perception. Motion direction discrimination was measured using random dot kinematograms (RDKs). Due to the increased distance between the stimulation and return electrodes in this montage, we expected a smaller reduction in motion discrimination thresholds compared to our previous study. The results suggest that increased interelectrode distance mitigates the efficacy of hf-tRNS. Additionally, no significant effects were observed with the other tES protocols tested. Our findings imply that the positioning of the two electrodes affects current flow characteristics, leading to reduced neuromodulation. These results underscore the importance of stimulation configuration, particularly the effect of interelectrode distance on performance. Given the widespread application of brain stimulation techniques in clinical and cognitive research, our results can guide future studies in carefully considering this further aspect of stimulation montage configurations.

## Introduction

In conventional electrical stimulation, both return and stimulation electrodes are placed over the scalp in different locations to alter cortical excitability within the reach of the electrical field generated by the current flow. Within this flow, application of an electric field to a neuron will instigate a change in the resting membrane potential, inducing an electric current to penetrate the membrane, which causes the cell to hyperpolarize or depolarize. This change in the membrane potential may reach the neuron’s threshold, triggering neuronal firing (Ye & Steiger, 2015). Alterations in neuronal population activity have been a promising method for modulating cortical excitability. Non-invasive techniques such as transcranial direct current stimulation (tDCS), where a weak direct current is applied to the scalp (Stagg & Nitsche, 2011), and more recently transcranial random noise stimulation (tRNS) wherein sinusoidal current alternates between two electrodes at a frequency spectrum (Antal et al., 2008), are powerful therapeutic tools. Reported effects of transcranial direct current stimulation (tDCS) and transcranial alternating current stimulation (tACS) include improvements in: motor function in stroke patients (Boggio et al., 2007), aphasia (Monti et al., 2008), depression (Nitsche et al., 2009; Moffa et al., 2020), phobia and anxiety (Cobb et al., 2021), epileptic seizures (Tekturk et al., 2016; Zoghi et al., 2016; San-Juan et al., 2017), schizophrenia (Cheng et al., 2020; Brunelin et al., 2021), tinnitus (Claes et al., 2014; Yuan et al., 2018; Mohsen et al., 2019), fibromyalgia (Curatolo et al., 2017), neuropathic pain in MS patients (Palm et al., 2016), and ADHD (Salehinejad et al., 2019; Berger et al., 2021). Compared to Sham, a-tDCS showed significant facilitation in visual working memory performance in patients with Alzheimer (Boggio et al., 2009). Conversely, tDCS is shown ineffective for generalized anxiety disorder (de Lima et al., 2019) and migraine (Pinto et al., 2018). While anodal tDCS (a-tDCS) demonstrated significant improvements in tinnitus symptoms (Fregni et al., 2006), cathodal tDCS (c-tDCS) was ineffective.

Random noise stimulation aims to deliver currents in a specific band of random frequencies, typically between 0-100 Hz for low-frequency tRNS (lf-tRNS) and 101-640 Hz for high-frequency tRNS (hf-tRNS) to alter cortical excitability. hf-tRNS was shown to induce long-lasting corticospinal excitability when applied to M1 (Terney et al., 2008; Potok et al., 2021). Cortical modulation due to tRNS further includes improvements in motion direction discrimination (Ghin et al., 2018), visual acuity (Moret et al., 2018), perceptual learning (Fertonani et al., 2011; Herpich et al., 2015), working memory (Murphy et al., 2020) and, improved perceptual learning in peripheral vision (Contemori et al., 2019). Further evidence has shown that tDCS and tACS can help alleviate migraine symptoms (O’Hare et al., 2021) and improve visual functions in myopia (Camilleri et al., 2016). Attenuated motion aftereffect (MAE) with lf-tRNS (0-100 Hz) and enhanced MAE modulation with hf-tRNS (101-640 Hz) have been reported (Campana et al., 2016), indicating independent functioning of low- and high-frequency tRNS (for a review on tRNS, see van der Groen et al., 2022). Moret et al. (2018) showed that hf-tRNS combined with perceptual training in amblyopia patients improved visual acuity but not contrast sensitivity thresholds.

On the other hand, many other tRNS studies have suggested no to mild effects. For instance, tRNS stimulation has been shown to be ineffective at improving outcomes in depression (Nikolin et al., 2020), and one session of tRNS did not show improvement in inhibitory control, whereas three sessions increased reaction time on a Go/No Go task (Brevet-Aeby et al., 2019). No improvement in error rates or reaction times were reported on a motor response inhibition task after tRNS (Brauer et al., 2018). Furthermore, occipital tRNS stimulation modulated sensitivity for low-contrast Gabor signal detection; however, it did not show an effect for high-contrast stimuli (Battaglini et al., 2019).

Electric field distribution has been of interest to researchers to identify the characteristics of current conducted throughout the brain tissue, which anatomically encompasses idiosyncratic components such as skin, skull, fibers, gyral folding, and cerebrospinal fluid (CSF), all of which affect electrical current flow (Asan et al., 2019; Grant & Lowery, 2009; Opitz et al., 2011). The electric field can also modulate other brain areas within the trajectory of the current flow and around the reference electrode, depending on the montages implemented (Parazzini et al., 2011; Opitz et al., 2016). Studies employing various electrode locations for electrical stimulation have led to somewhat contradictory outcomes. For instance, Moliadze et al. (2010) observed changes in motor-evoked potentials (MEPs) with extended return electrode distance using anodal tDCS (a-tDCS) and tRNS. When stimulating at 1 mA they observed increased cortical excitability only with cranial stimulation; however, increasing the amplitude to 2 mA induced excitability in both cranial and extracranial configurations. In other words, the loss of excitability was compensated by increased stimulation intensities. Moreover, a loss of cortical excitability was again observed when the distance was further increased.

Additionally, repetitive tDCS with an extracephalic electrode configuration showed greater improvements in major depressive disorder compared to a bifrontal configuration (Martin et al., 2011). In a simulation study using Finite Element Method (FEM), Noetscher et al. (2014) suggested that extracephalic montage might increase total current densities in white matter, induce larger average vertical current densities in the primary motor cortex and somatosensory cortex, and cause no change or smaller averages in horizontal current densities for specific cortical areas compared to a cephalic montage. Furthermore, it has been shown that cathodal tDCS (c-tDCS) with an extracephalic return electrode arrangement impairs cognitive inhibition (Friehs & Frings, 2019). Accornero et al. (2007) measured visual evoked potentials (VEPs) using tDCS, placing the return electrode at the base of the neck. They reported decreased VEPs with a-tDCS, but an increased amplitude of the P1 component with c-tDCS was observed.

Fertonani et al. (2011) tested visual perceptual learning performance while employing different stimulation types, such as a-tDCS, c-tDCS, low- or high-frequency tRNS (hf-tRNS), or Sham stimulation, to investigate potential effects on behavioral performance. The stimulation electrode was placed over V1, whereas the return electrode was positioned extracephalically on the right arm. Among all the stimulation types, hf-tRNS was shown to yield superior results in improving visual perceptual learning. Using an extracephalic electrode and targeting the ventromedial prefrontal cortex (VMPFC) with c-tDCS Yin et al. (2021) showed that the self-prioritization effect was abolished. In another study somewhat mixed results were observed in children with ADHD: no improvements in working memory, cognitive flexibility, or response inhibition were seen with either a-tDCS or c-tDCS, whereas only unilateral a-tDCS over the dorsolateral prefrontal cortex (DLPFC) showed partial improvements in executive control (Salehinejad et al., 2022). Both a-tDCS and c-tDCS resulted in improved accuracy and reaction time regarding executive function, while decreased working memory performance with c-tDCS was observed in active-duty soldiers (Duffy et al., 2024).

In sum, contradictory results have been shown regarding transcranial electrical stimulation (tES) efficacy, which may be due to factors such as experimental setup, behavioral paradigms, stimulation magnitudes, electrode locations, and interindividual variability (Rodella et al., 2021). Electrode configuration appears to be one of the crucial factors contributing to these inconsistencies. To our knowledge, no investigation has yet focused on comparing the efficacy of cephalic versus extracephalic electrode placements using a motion discrimination paradigm and examining the effects of different stimulation techniques, such as a-tDCS, c-tDCS, and hf-tRNS. Here, we aimed to address design-related disparities in the literature by combining different stimulation types and electrode arrangements within the same behavioral paradigm, thereby presenting more integrated results.

Ghin et al. (2018) employed a motion coherence discrimination task using anodal or cathodal tDCS, hf-tRNS, or Sham stimulation. They introduced a cephalic electrode montage, placing the active electrode on the left hMT^+^ and the return electrode over the vertex (corresponding to the Cz electrode on a 64-channel EEG system). They found reduced motion coherence thresholds only when the stimulus was presented in the contralateral visual hemifield with respect to the stimulated hMT^+^. In this study, we investigated the effects of tDCS and hf-tRNS on motion discrimination performance, with particular emphasis on electrode configuration (i.e.: distance) across different brain stimulation types. We expected diminished effects of tES as the distance between the stimulation and return electrodes increased, as suggested by Moliadze et al. (2010) and Opitz et al. (2016). Therefore, the motion coherence threshold alteration induced by hf-tRNS with a cephalic montage, as shown in Ghin et al. (2018), should be reduced or abolished with an extracephalic arrangement. Furthermore, we asked whether other tES types may have any effect on motion coherence thresholds using the same extracephalic arrangement. We compared the cephalic electrical stimulation data from Ghin et al. (2018) with the current extracephalic stimulation data to explore the effects of interelectrode distance.

## Methods

The methods used in this experiment were adapted from Ghin et al. (2018). The paradigm employed in their first experiment was implemented to investigate the effects of inter-electrode distance.

### Participants

One of the authors (AP) and twelve naïve participants took part in the experiment. All participants had normal or corrected-to-normal vision. All experimental protocols and procedures were in accordance with the World Medical Association Declaration of Helsinki (2013) and approved by the Ethics Committee of the University of Lincoln. All participants were screened with a questionnaire for implanted metal objects, any past or present history of seizures, heart-related problems, or any neurological issues to be excluded from the study. They received detailed information about the protocol and signed a consent form. Participants were given monetary compensation upon their participation in the experiment. In the current study, a post hoc Power analysis was conducted using G*Power (3.1) (Faul et al., 2007, 2009) to determine the statistical power achieved with our sample size. The total sample size of 28 was derived from the sum of the sample sizes of two experiments: the cephalic montage experiment (Ghin et al., 2018) with 16 participants and the extracephalic montage experiment with 13 participants. It should be noted that one participant (RD) who took part in both experiments was excluded from the cephalic montage experiment to maintain independence of observations, hence the total of 28 participants. The power analysis was based on the partialrJ^2^ values obtained from our mixed ANOVA, which were converted to Cohen’s *f* effect size. With an alpha level set at 0.05, eight measurements and two groups, the Power analysis indicated that our study achieved a power of approximately 0.95. This suggests that our study was sufficiently powered to detect large effects of interest in the interaction between visual field, stimulation type, and experimental group.

### Apparatus

Stimuli were presented on a 20-inch HPp1230 monitor with a refresh rate set to 85 Hz. The screen resolution of the monitor was 1280×1024 pixels, with each pixel corresponding to 1.6 arcmins. The minimum and maximum luminance values of the screen were 0.08 and 74.6 cd/m^2^ respectively, and the mean luminance was 37.5 cd/m^2^. Stimuli were displayed on a gamma-corrected monitor using a look-up table (LUT) to linearize the luminance range. MATLAB Psychtoolbox-3 was used to generate stimuli and data collection (Brainard, 1997; Pelli, 1997; Kleiner, Brainard, & Pelli 2007). Participants were seated at a 57 cm distance from the screen using a chin and forehead rest.

### Stimuli

Stimuli were random dot kinematograms (RDKs) made of 150 randomly moving white dots (0.12 deg diameter). RDKs were presented within a circular aperture with a diameter of 8 deg (density: 3 dots/deg^2^). Dot drifting speed was 13.4 deg/s, and the eccentricity was 12 deg. However, for four participants the task with the speed of 13.4 deg/s was too difficult (occasionally producing perception of opposite motion direction; Bae & Luck, 2022). For these participants only, we used a speed of 8 deg/s. Each dot had a limited lifetime of 47 ms before it disappeared and reappeared in another random location within the circular aperture. Dot density was maintained by replacing the dots that moved out of the circular window, repositioning them randomly at another location. A proportion of the dots moved in a coherent direction (signal dots), while the remaining moved in random directions (noise dots). The probability of a dot to be signal or noise was kept constant and the dot appearance on the screen was asynchronous (Morgan & Ward, 1980; Newsome & Pare, 1988; Geisler, 1999). RDKs were displayed for approximately 106 ms and appeared either on the right or the left side of the fixation point, randomly selected and counterbalanced.

### Procedure

Participants were asked to fixate their gaze on the fixation point and respond to the direction of the RDKs presented on either visual hemifield, i.e., right or left to the fixation point (see Fig.1 in Ghin et al. (2018) for a schematic representation of the stimulus and the procedure section for more details). Participants performed an eight alternative forced choice (8AFC) task with a key response. We estimated the motion coherence thresholds that corresponded to 70.7% accuracy.

**Figure 1.**
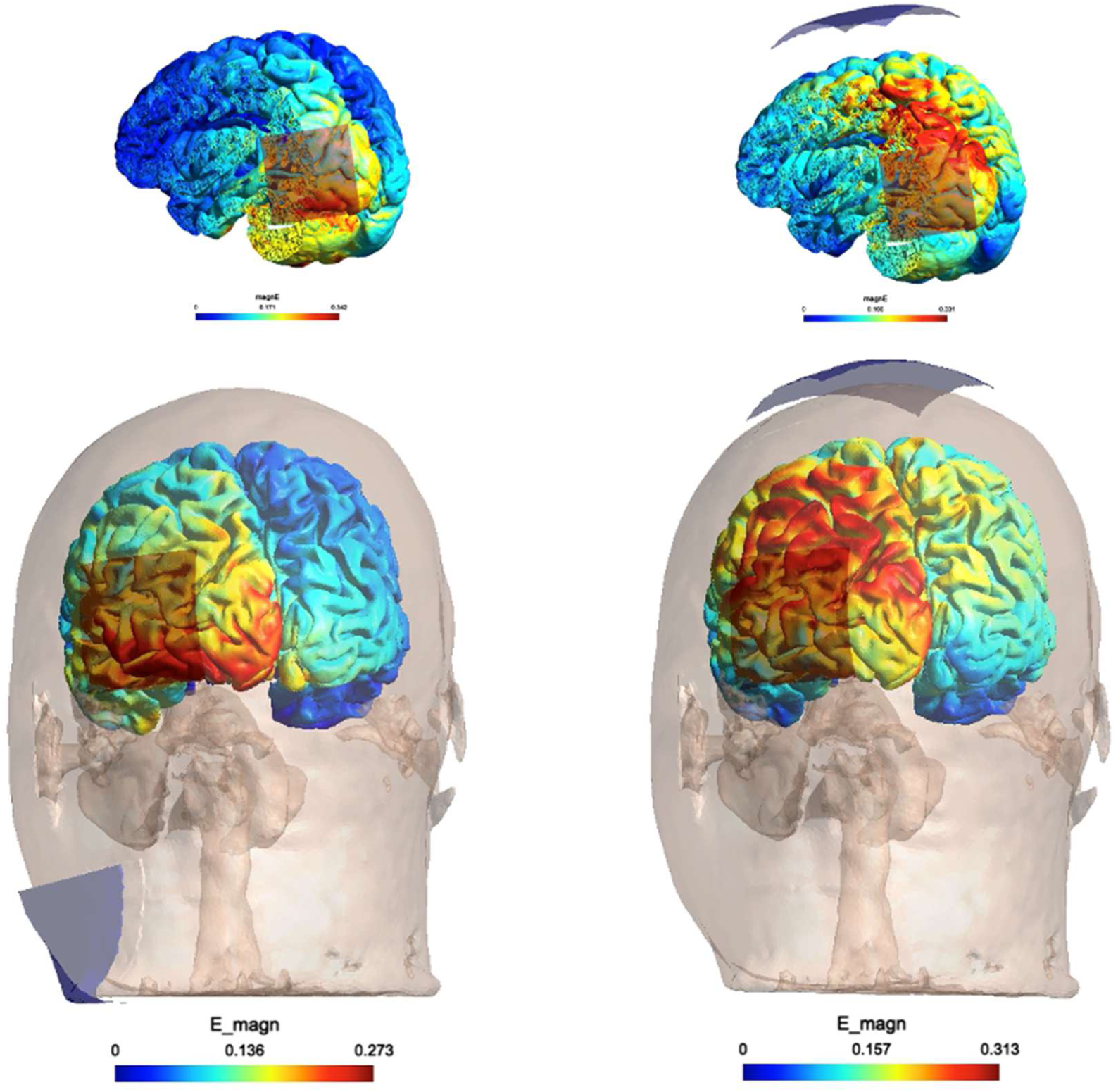
tES EF modeling corresponds to the extracephalic montage (left panel) and to the cephalic montage from Ghin et al. (2018) (right panel, rerun with SimNIBS). Electrode locations: PO7-neck location for extracephalic montage, PO7-Cz for the cephalic montage.

Coherence thresholds and slope of the psychometric function were estimated based on the observances from five interleaved adaptive staircases (MLP) (Grassi & Soranzo, 2009; Green, 1993). One block consisted of 5 staircases for each visual hemifield, and each staircase consisted of 32 trials. All four stimulation types (a-tDCS, c-tDCS, hf-tRNS, and Sham stimulation) were delivered throughout the task on separate, non-consecutive days, i.e., one session each day, and each session consisted of five blocks. Participants were not informed of the type of stimulation used for the sessions.

### Electrical Stimulation

A BrainSTIM EMS device (http://www.brainstim.it/index.php?lang=en) was utilized for electrical stimulation. The intensity of stimulation was 1.5 mA with a 30 second ramp up. Alternating current oscillating at random frequencies within the range of 100-600 Hz was delivered for the hf-tRNS condition. Sham condition consisted of 30 seconds of stimulation at the beginning of the trial, but no electrical stimulation was delivered during the task (Gandiga et al., 2006). Stimulation and return electrodes were 16 cm^2^ and 60 cm^2^, respectively. Stimulation safety was within the safety criteria described before (Poreisz al., 2007; Bikson et al., 2016). One electrode was placed on hMT^+^, the area responsible for visual motion processing (Albright, 1984; Newsome & Pare, 1988; Maunsell & Van essen, 1983; Tootell et al., 1995; Smith et al., 2006), and the other electrode on the ipsilateral deltoid muscle to devise an extracephalic configuration (Fertonani et al., 2011). Since the stimulation electrode was placed over the left hMT^+^, the left visual field remained ipsilateral to the stimulated area. Therefore, the contralateral (right) visual hemifield was expected to show the dominant modulation on motion discrimination thresholds and the slope.

Approximately 18 minutes of stimulation was employed on each session, and the 4 stimulation sessions (one for each stimulation type) were conducted on different days. The target area was localized in all observers by using predetermined coordinates: 3 cm dorsal to inion and 5 cm leftward from there for the localization of the hMT^+^ (Campana et al., 2006; Campana et al., 2002; Campana et al., 2013; Laycock et al., 2007; Pascual-Leone et al., 1999; Pavan et al., 2011, 2017; Battelli et al., 2002; Edwards et al., 2017; Théoret et al., 2002); this site provides a localization that is consistent with fMRI localizers (Thompson et al., 2009).

### Estimated Electric Fields Induced by tES

Simulations on SimNIBS 4.1 were performed (Thielscher et al., 2015; Saturnino et al., 2019) to estimate the potential electric field distribution over hMT^+^. An extracephalic configuration was employed with one electrode over the area corresponding to hMT^+^ as described in the methods section and the other electrode on the ipsilateral neck location given the spatial limitations of the mesh used in the software that extends to the neck without the shoulder. Regarding the return location, Maas et al. (2023) showed that using SimNIBS, electric field (EF) estimates were largely consistent using lower neck or shoulder return locations. Van Hoornweder et al. (2024) suggested neck vs shoulder simulations only showed a 0.011 V/m peak magnitude difference. Electrode sizes and sponge measurements were kept the same as in the verum tDCS stimulation. Extracephalic and cephalic electrode configurations were simulated to assess electric field distribution, following the parameters set by Ghin et al. (2018). The simulations revealed that the 99.9^th^ percentile of the electric field for the extracephalic montage was 0.307 V/m, slightly lower than the 0.328 V/m observed for the cephalic configuration (Figure 1). Furthermore, focality analysis indicated that 75% of the 99.9^th^ percentile field was confined to a volume of 1.15 · 10⁴ mm^3^ in the extracephalic montage, compared to 1.45 · 10⁴ mm^3^ in the cephalic configuration. These results suggest that while both configurations produce similar peak electric field magnitudes, the cephalic montage results in a slightly larger, more diffuse field distribution compared to the more focused field generated by the extracephalic setup.

In order to extrapolate the above results to the case of alternating current, we used a convenient approximation that was first proposed in Ghin et al. (2018). Such approximation allows to extrapolate the expected values of the peak electric field magnitude generated by an alternating current stimulus of given intensity, in terms of the electric field generated by a direct current of equal intensity, if the frequency of the stimulus lies below a threshold value that was quantified in Ghin et al. (2018) in the order of kHz.

The idea underlying the approximation is that the electric field E_ω_ generated by an alternating current stimulus of frequency ω is proportional to the one E_O_ generated by a direct current (that is, when ω → 0) via a scaling factor *r* defined as:

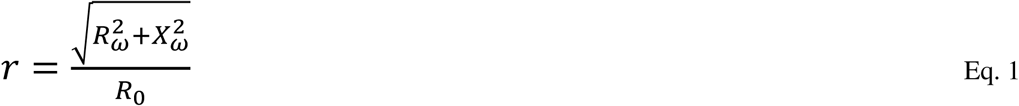

*R* and *X* being the real and imaginary components of the electrical impedance across the path between the two electrodes (the subscripts refer to the ω frequency stimulus and the direct current stimulus, respectively). Therefore, the knowledge of these components allows us to compute E_ω_ from E_0_.

Unfortunately, the impedance values used in Ghin et al. (2018) refer to a shorter, intracephalic path, and we were not able to retrieve from the literature any estimate of these values for the longer path considered in the present study. Nonetheless, if we make the further assumption that the impedance grows linearly with the path length (which is formally true only along a perfectly homogeneous path, but is still an acceptable approximation as long as several yet similar body tissues are crossed), both the numerator and the denominator of Equation (1) are increased by the same (path length-related) factor, resulting in the same ratio *r* as in the previous case.

Taking all the above considerations into account, it is possible to claim that the peak electric field experienced in the extracephalic montage ranged approximately from 0.146 V/m at ω = 100 Hz (when *r* was 0.476) to 0.054 V/m at ω = 600 Hz (when *r* was 0.177) whereas the cephalic montage ranged approximately from 0.156 V/m at ω = 100 Hz (when *r* was 0.476) to 0.058 V/m at ω = 600 Hz (when *r* was 0.177). It is important to note that these values are based on a series of approximations and should be regarded as estimates serving only the purpose of this consistency check.

### Data analysis

To test for differences in coherence threshold between subgroups of participants exposed to the stimulus at different speeds, we used a permutation ANOVA (10,000 permutations) with global motion speed as the between-subjects factor. Subsequently, data from the two groups (extracephalic and cephalic montage groups) were analyzed using a three-way mixed ANOVA with one between-subjects factor (the group/montage) and two within-subjects factors (visual hemifield and stimulation type). Post hoc comparisons were corrected with the FDR method (α = .05). The normality of residuals was assessed using QQ plots and a Shapiro-Wilk test.

The skewness of residual distributions was also assessed. Outliers were identified using the boxplot method, where values above Q3 + 1.5IQR or below Q1 - 1.5*IQR were considered as outliers. Q1 and Q3 are the first and third quartile, respectively, while the interquartile range (IQR) is defined as IQR = Q3 - Q1. The study focused on two dependent variables: the coherence thresholds corresponding to 70.7% correct discrimination and the slope of the psychometric function. Details of the staircase operational workflow and the calculations for determining coherence threshold and psychometric function slope are provided in Appendix A. All analyses and visualizations were conducted using MATLAB 2022b, *R* (v4.3.3) software (R Core Team, 2023) and *Jamovi* 2.4.

The materials, data, and scripts used in these experiments are available on the Open Science Framework (OSF) at the following link: https://osf.io/tdq3p/

## Results

### Coherence Thresholds

#### Permutation ANOVA

The permutation ANOVA was performed using the ‘*perm.anova*’ routine from the ‘*RVAideMemoire*’ R package (Hervé, 2020). The results showed no significant difference between the two subgroups in the extracephalic montage when using global motion at different speeds (*F*1 = .0032, *p* = .952).

### Omnibus three-way mixed ANOVA

Figure 2 shows the coherence thresholds for each hemifield and stimulation condition, separately for the cephalic and extracephalic montage. From the QQ plots and the Shapiro-Wilk tests performed for all the conditions, residuals were approximately normally distributed (all *p*s > .05). The skewness was around 1 or lower across all conditions, indicating small or negligible deviance of data distribution from symmetry. The assumption of homogeneity of variances was also met. We identified one outlier using the boxplot method that was included in the analysis (cephalic montage group, left visual hemifield, c-tDCS, coherence threshold = 68.1).

**Figure 2.**
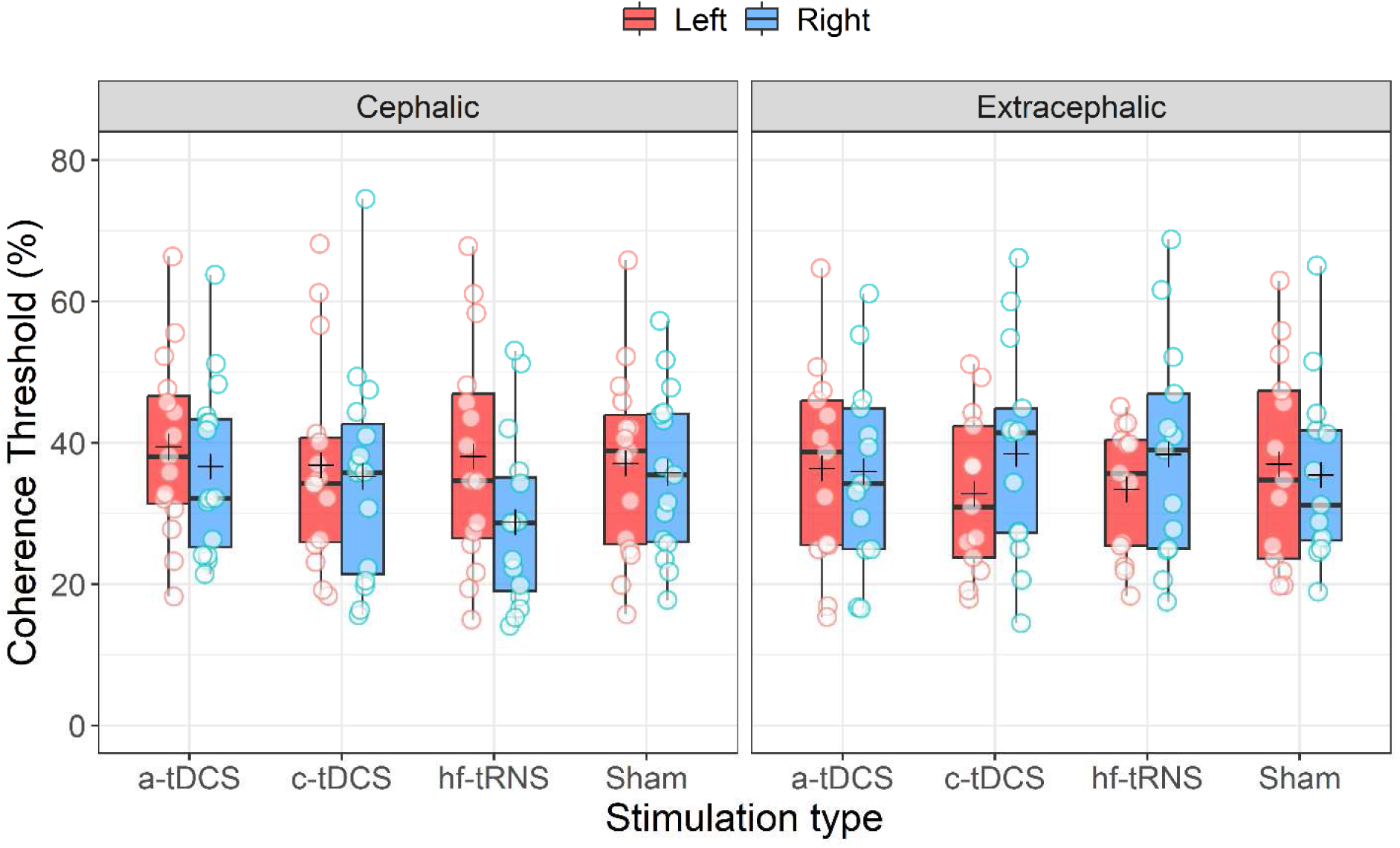
Boxplots of coherence thresholds by stimulation type, categorized by the montage type (cephalic and extracephalic) and visual hemifield. The black thick horizontal line inside each box is the median, whereas the cross represents the mean.

The omnibus mixed ANOVA revealed only a significant interaction between group and visual hemifield (*F*1, 26 = 7.27, *p* = .012, p2=.22) and the three-way interaction between group, visual hemifield, and stimulation type (*F*3, 78 = 3.05, *p* = .034, p2=.11). All the other main effects and interactions were not significant (group/montage: *F*1, 26 < .001, *p* = .99, p2<.001; visual hemifield: *F*1, 26 = .52, *p* = .48, p2=.02; stimulation: *F*3, 78 = .80, *p* = .5, p2=.03; stimulation x group: *F*3, 78 = .62, *p* = .61, p2=.023; visual hemifield x stimulation: *F*3, 78 = 1.09, *p* = .36, p2=.04). For the interaction between group and visual hemifield, FDR corrected post hoc comparisons (correction applied for six comparisons) did not reveal any significant difference (all *padj* > .05).

For the three-way interaction, we performed a two-way repeated measures ANOVA at each level of the group/montage. Specifically, one two-way RM ANOVA for the extracephalic montage and another for the cephalic montage.

For the extracephalic montage, the two-way RM ANOVA did not reveal any significant main effect or interaction (visual hemifield: *F*1, 12 = 1.85, *p* = .2, p2=.134; stimulation: *F*3, 36 = .024, *p* = .99, p2=.002; visual hemifield x stimulation: *F*3, 36 = 1.51, *p* = .23, p2=.11).

For the cephalic montage, the two-way RM ANOVA revealed a significant effect of the visual hemi-filed (*F*1, 14 = 6.20, *p* = .026, p2=.31) and a significant interaction between visual hemifield and stimulation (*F*3, 42 = 2.9, *p* = .046, p2=.17). The main effect of stimulation was not significant (*F*1.83, 25.63 = 1.64, *p* = .21, p2=.11). For the stimulation factor, the sphericity was violated; therefore, we used the Greenhouse-Geisser correction for the degrees of freedom. For the visual hemifield, on average, coherence thresholds were lower for the right visual hemifield (mean difference: 3.73%; SE: 1.50%). For the visual hemifield x stimulation interaction, FDR-corrected post hoc (correction applied for 28 comparisons) showed a significant difference between left hemifield a-tDCS and right hemifield hf-tRNS conditions (*padj* = .0448), between left hemifield c-tDCS and right hemifield hf-tRNS conditions (*padj* = .0009), between left and right hemifields in the hf-tRNS condition (*padj* = .0009), the latter with a mean difference of 9.25% (SE: 1.91%). A significant difference was also found between left Sham and right hf-tRNS (*padj* = .0009) and between right Sham and right hf-tRNS conditions (*padj* = .0448). All the other comparisons did not reach the significance level. These results suggest that hf-tRNS facilitates global motion discrimination by lowering the coherence threshold in the contralateral visual field, but only with a cephalic montage, i.e., when both electrodes are placed on the scalp (see Figure 2).

### Slope

Figure 3 shows the slopes for each hemifield and stimulation condition, separately for the cephalic and extracephalic montage. The permutation ANOVA on the slope showed no significant difference between the two subgroups with different global motion speeds in the extracephalic montage (*F*1 = .9709, *p* = .335).

**Figure 3.**
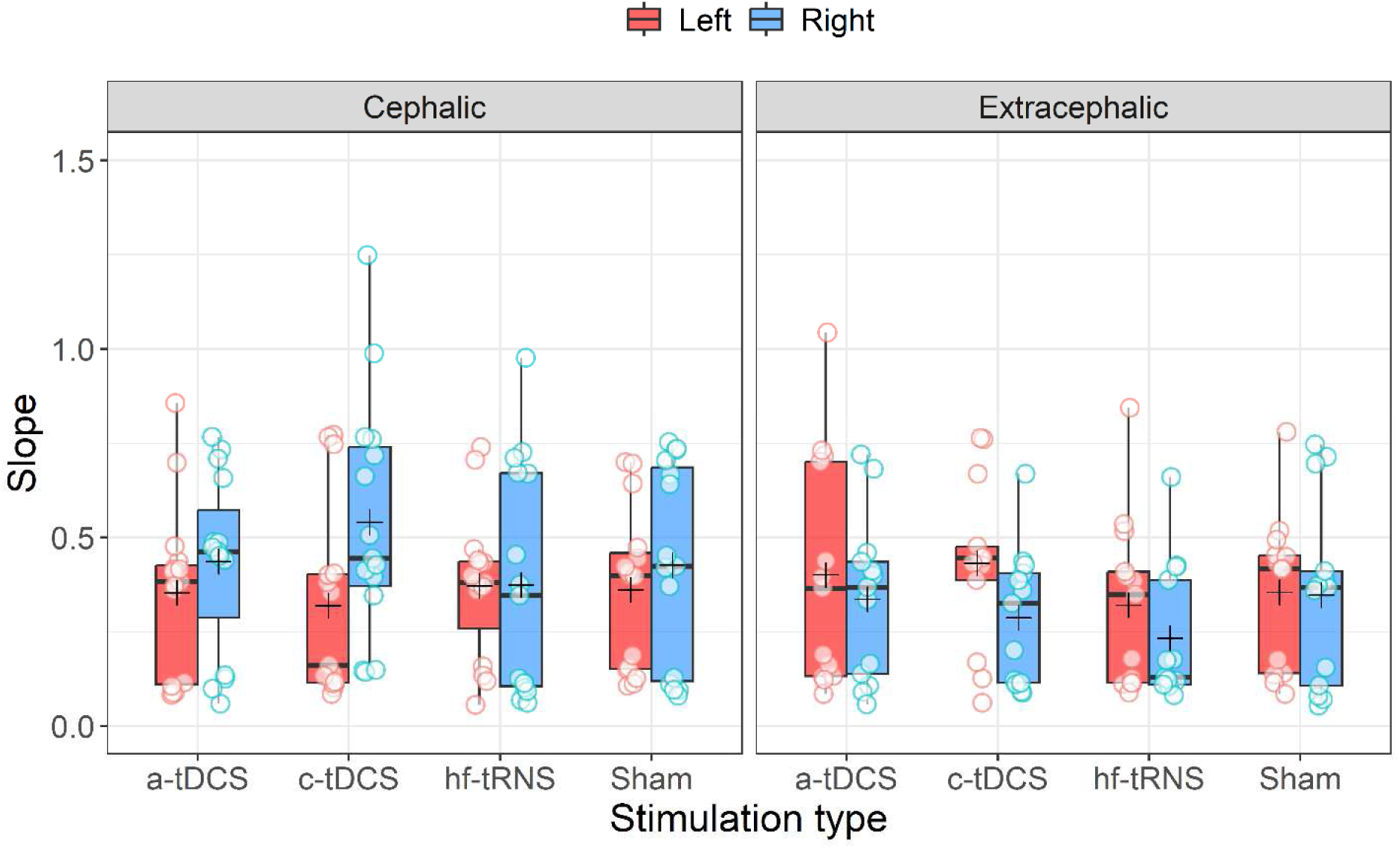
Boxplots of slopes by stimulation type, categorized by the montage type (cephalic and extracephalic) and visual hemifield. The black thick horizontal line inside each box is the median, whereas the cross represents the mean.

From the QQ plots and the Shapiro-Wilk tests performed for all the conditions split by the two groups/montage types, residuals were approximately normally distributed but in four conditions (cephalic, left visual field, a-tDCS, *p* = .049; cephalic, left visual field, c-tDCS, *p* = .003; cephalic, right visual field, hf-tRNS, *p* = .022; extracephalic, right visual field, hf-tRNS, *p* = .003; cephalic, right visual field, Sham, *p* = .018). However, the skewness was always < 1 suggesting not much deviation from symmetry. The assumption of sphericity and homogeneity of variances were met. We identified two outliers with the boxplot method that were included in the analysis. The omnibus mixed ANOVA revealed only a significant interaction between group and visual hemifield (*F*1, 26 = 5.44, *p* = .028, p2=.17). All the other main effects and interactions were not significant (group/montage: *F*1, 26 = 3.48, *p* = .074, p2=.12; visual hemifield: *F*1, 26 = .06, *p* = .81, p2=.002; stimulation: *F*3, 78 = .84, *p* = .48, p2=.031; stimulation x group: *F*3, 78 = .21, *p* = .89, p2=.008; visual hemifield x stimulation: *F*3, 78 = .36, *p* = .79, p2=.013; visual hemifield x stimulation x group: *F*3, 78 = 1.213, *p* = .31, p2=.045). For the interaction between group and visual hemifield, FDR-corrected post hoc comparisons (correction applied for six comparisons) did not reveal any significant difference (all *padj* > .05). These results suggest that the tES techniques applied had no effect on the slope, which is related to stimulus discriminability.

## Discussion

Differences in the literature regarding clinical or cognitive applications of brain stimulation were linked to parameters such as electrode size, position, stimulation type and intensity, duration, and frequency. The extracephalic configuration was preferred when targeting deeper regions (Noetscher et al., 2014, Bai et al., 2014; Guidetti et al., 2023) or to avoid stimulating additional areas under the return electrode. In this study, we aimed to investigate how increasing interelectrode distance, as it occurs with extracephalic montage, has significant consequences in producing modulatory effects.

Our results showed that extracephalic conditions in general did not induce changes on motion coherence thresholds: no stimulation type altered performance compared to Sham conditions. However, in the cephalic montage, where the interelectrode distance is much smaller, the effect of stimulation was observed between right and left hemifield conditions. The hf-tRNS condition in the cephalic montage resulted in strong modulations when the global moving RDK was presented in the visual hemifield contralateral to the site of stimulation. Coherence thresholds were not only lower for the right visual hemifield in the hf-tRNS condition when compared to the right visual hemifield Sham condition, but also to the left visual hemifield across the hf-tRNS, a-tDCS, c-tDCS, and Sham conditions. In other words, stimulating over left hMT^+^ with hf-tRNS overall improved performance (i.e., lowered coherence thresholds) on the motion direction discrimination task presented in the contralateral hemifield.

In this study we run simulations for both cephalic and extracephalic montages to compare the estimated EF distributions. Figure 1 shows the results of increasing the interelectrode distance. Traditionally, when a specific region is intended to be modulated by electrical stimulation, a stimulation electrode is placed over that area, expecting to induce cortical excitability changes. However, while the cephalic montage estimations appear to target areas responsible for motion perception, such as hMT^+^, extracephalic montage seems to shift the field away from it towards regions such as the inferior occipital gyrus (IOG), the posterior inferior temporal gyrus (pITG), the temporal pole (TP; not shown Figure 1), and the cerebellum. These areas are associated with functions such as face selectivity (IOG; Jacques et al., 2019; de Haas et al., 2021), semantic control (pITG; Hodgson, 2023), face recognition and semantic processing (TP; Herlin et al., 2021), object comprehension in general (the whole anterior temporal lobe; Bonner & Price, 2013), and movement (cerebellum), none of which directly relate to motion processing. This shift may account for the lack of modulation in coherence thresholds observed across all stimulation types (a-tDCS, c-tDCS, hf-tRNS) in the extracephalic configuration.

To investigate the simulation profiles of few studies which used extracephalic montage and reported lack of behavioral improvements we conducted SimNIBS simulations to explore the EF distribution on targeted areas. Salehinejad et al. (2022), placing the stimulation electrode over the dorsolateral prefrontal cortex (DLFPC), showed no improvement on the executive functioning task performance using tDCS. Running a simulation of their stimulation parameters reported in their paper showed the presence of higher EFs on the ventrolateral prefrontal cortex (VLPFC) rather than their initial target area, potentially contributing to their behavioral results. Simulation results for Duffy et al. (2024) showed high EFs over VLPFC when the stimulation electrode was positioned over the DLPFC. Their results were mixed, with increased reaction time and d′ for males with a-tDCS and females with c-tDCS. Fertonani et al. (2011) showed modulation of d′ with hf-tRNS compared to Sham, a-tDCS, c-tDCS. Running their stimulation parameters revealed that both occipital lobes remained within the peak EF distribution, making it reasonable to observe corresponding behavioral effects. Martin et al. (2011) showed that repetitively applying extracephalic tDCS on DLPFC improved depression scores more than bifrontal stimulation, likely due to the widespread EF distribution across the brain, including the limbic areas associated with symptoms of depression.

In addition to adjusting the interelectrode distance, reducing focality could decrease targeting of hMT^+^, as previous research has shown that focality decreases with increased distance in a cephalic montage (Mackenbach et al., 2020; Nitsche et al., 2007). Other ways of improving focality could be to decrease the electrode size (Faria et al., 2011), use a ring electrode (Datta et al., 2009) or increase the return electrode size (Nitsche et al., 2007). Increased intensity could also potentially influence the behavioral output as suggested in Moliadze et al. (2010) however, estimated spatial distribution of EFs should be considered while manipulating the stimulation intensity.

This study aimed to present a general overview of how electrode distance contributes to behavior as a result of electrical stimulation. Based on the simulations, targeting the region of interest below the stimulation electrode seems more precise with a smaller inter-electrode distance whereas, in contrast, increasing the distance by extending one electrode to a further location may have unwanted consequences. Therefore, inter-electrode distance should carefully be planned out while implementing a tES protocol.

Optimizing tES protocols remains a critical challenge considering its widespread clinical use in today’s therapeutic applications. Promising tES outcomes in the literature points to an extensive therapeutic potential. Therefore, stimulation parameters such as electrode size, interelectrode distance, stimulation type and intensity need to be carefully addressed, and application protocols must be tailored according to the desired therapeutic outcomes.

## Data availability

The materials, data, and scripts used in these experiments are accessible on the Open Science Framework (OSF) at this link: https://osf.io/tdq3p/

## Acknowledgments

This study was supported by the College of Social Science of the University of Lincoln.

## Contributions

**A.P.** conceived and designed the experiment. **A.P.** implemented the experiment. **A.P., S.K.Y.**, and **J.F.** collected the data. **A.P., S.K.Y., H.K.**, and **J.F.** analyzed data. **A.P., J.F., S.K.Y., H.K.**, and **M.W.G.** interpreted the results. **A.P., J.F., S.K.Y., H.K.**, and **M.W.G.** wrote the main manuscript. All authors reviewed the manuscript.

## Appendix A Estimation of coherence threshold and slope from MLP

The operational flow of the staircase to estimate coherence threshold and slope of the psychometric function consisted in acquiring and storing the subject response to the *n*-th trial, selecting the psychometric function maximizing the likelihood of the first *n* trials, estimating the corresponding coherence threshold, and presenting it as stimulus for the (*n*+1)-th trial. The estimate subsequent to the last trial was the output of the staircase (Grassi & Soranzo, 2009). The logistic function was used as psychometric function:

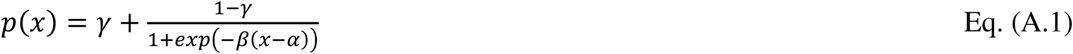

The baseline parameter *γ* was fixed to 1/8, while the midpoint *α* and the slope parameter *β* were varied to maximize the likelihood. The rationale for such a choice was to focus on the position of the threshold on the coherence axis, suppressing the further degree of freedom associated with the growth rate of the psychometric function. However, for the sake of completeness, we also extracted information about the slope. To do this, we made use of a custom best fit routine based on a Metropolis-Hastings algorithm, exploring the parameter space of the logistic function. The algorithm randomly selected a starting point in the parameter space {*α, β*} and computed the corresponding total likelihood:

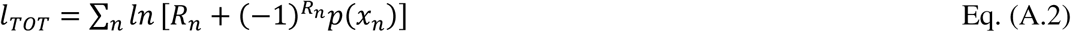

over the whole staircase. Here *x_n_* is the coherence of the *n*-th trial, while *R_n_* indicates the corresponding subject response (1 for correct, 0 for wrong). Thereafter, during each iteration of the Metropolis-Hastings, it performed a random step in the parameter space, computed the corresponding total likelihood and compared it to the one of the starting points. If the new likelihood was higher, the algorithm replaced the starting point with the new point, thus accepting the step. Otherwise, the step was rejected. Approximately 100,000 iterations were performed for each staircase, and the logistic function corresponding to the highest likelihood was returned as the best fitting curve. Using the best fit parameters, it was possible to compute an estimate for the coherence threshold *T_c_* as the inverse logistic function:

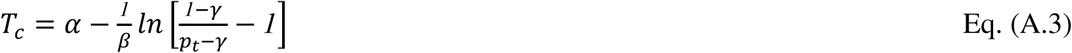

*p_t_* being the 70.7% accuracy value acquired by the psychometric function in correspondence of the coherence threshold. The slope of the psychometric function was calculated as:

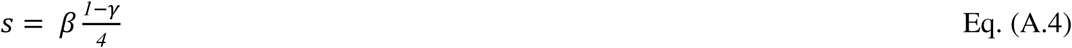

where y = .*125*.

